# Improved sgRNA design in bacteria via genome-wide activity profiling

**DOI:** 10.1101/272377

**Authors:** Jiahui Guo, Tianmin Wang, Changge Guan, Bing Liu, Cheng Luo, Zhen Xie, Chong Zhang, Xin-Hui Xing

## Abstract

CRISPR/Cas9 is a promising tool in prokaryotic genome engineering, but its success is limited by the widely varying on-target activity of single guide RNAs (sgRNAs). Based on the association of CRISPR/Cas9-induced DNA cleavage with cellular lethality, we systematically profiled sgRNA activity by co-expressing a genome-scale library (~70,000 sgRNAs) with Cas9 or its specificity-improved mutant in *E. coli*. Based on this large-scale dataset, we constructed a comprehensive and high-density sgRNA activity map, which enables selecting highly active sgRNAs for any locus across the genome in this model organism. We also identified ‘resistant’ genomic loci with respect to CRISPR/Cas9 activity, notwithstanding the highly accessible DNA in bacterial cells. Moreover, we found that previous sgRNA activity prediction models that were trained on mammalian cell datasets were inadequate when coping with our results, highlighting the key limitations and biases of previous models. We hence developed an integrated algorithm to accurately predict highly effective sgRNAs, aiming to facilitate the design of CRISPR/Cas9-based genome engineering or screenings in bacteria. We also isolated the important sgRNA features that contribute to DNA cleavage and characterized their key differences among wild type Cas9 and its mutant, shedding light on the biophysical mechanisms of the CRISPR/Cas9 system.

---

Efficient and reliable genome editing tools play crucial roles in genome engineering of prokaryotic hosts (1–8). The recently reported CRISPR/Cas9 system exhibits several advantages as a novel genome editing tool (9, 10). The system consists of a nuclease activity–carrying Cas9 protein and specificity-programming single guide RNA (sgRNA), the latter of which targets the complex to a genomic region flanked by a 3’NGG protospacer adjacent motif (PAM) via Watson-Crick base pairing (11). It works by introducing a double-strand break (DSB) in the chromosome, which is lethal to many prokaryotic hosts. The DSB then serves as a selection pressure to enrich for mutations introduced via homologous recombination with the artificial donor DNA. This method is broadly applicable to many prokaryotic organisms (10, 12–15), especially archaea (16). It also enables multiplex genome editing in a marker-free manner (17, 18), saving substantial time and labor during genome engineering. Lastly, only ~20 nucleotides in sgRNA encode the target of CRISPR/Cas system, compatible with massively parallel microarray oligonucleotide synthesis and next generation sequencing (NGS), both of which simplify the procedure for performing large-scale engineering or functional genomics studies (8, 19, 20).

The success of the CRISPR/Cas9 system for genome engineering of prokaryotic hosts is largely based on the activity of the selected sgRNA, or namely the cellular lethality caused by CRISPR/Cas9 as guided via a particular sgRNA to target the locus of interest. Poor sgRNA activities result in a high rate of false positives during genome editing, which results in the survival of many wild-type cells within the population. Conventional belief holds that DNA in prokaryotic cells is less protected than that in eukaryotic cells because of the lack of complex chromatin structures (21), and thus genome editing systems should typically work well in prokaryotic organisms. Studies, however, have suggested otherwise the existence of inactive sgRNAs during genome editing in bacterial cells (8, 13, 22, 23). This problem is especially prevalent when CRISPR/Cas9 genome editing is used in a multiplex manner, a major proposed advantage of this system, as the percentage of successfully modified cells decreases exponentially when sgRNA activity is not optimized (17). Equally deleterious when CRISPR/Cas9 is used for pooled screening within prokaryotic functional genomics studies, is that the diversity of sgRNA activities introduces noise in the downstream data processing to identify the mutations responsible for the phenotype under investigation (8).

Despite our lack of knowledge about the mechanisms responsible for sgRNAs with poor activity and their impact on the successful application of CRISPR/Cas9 genome editing in prokaryotic hosts, to the best of our knowledge, no investigation has yet been performed to systematically address this issue. By contrast, several recent pioneering studies have described the sgRNA sequence-activity relationship and resulted in corresponding prediction algorithms based on experimentally produced large datasets from mammalian cell lines (24–29). It is, however, worth noting that key differences exist between eukaryotic and prokaryotic hosts for CRISPR/Cas9-based genome editing. Crucially, mammalian cells have a highly active non-homologous end-joining (NHEJ) pathway (30), which plays fundamental roles in CRISPR/Cas9-induced DSB repair via an error-prone manner (20), rendering the reported dataset of eukaryotic sgRNA activity (25–27) a hybrid output that combines the inherent features of the sgRNAs with the known NHEJ preference for different DSB substrates (31). In addition, the complex chromatin structure that is unique to eukaryotic chromosomes, specifically, the blocking effect of nucleosome is paramount to reshape the sgRNA activity landscape (26, 32). It is thus reasonable to be skeptical of the general applicability of the established conclusions from studies in eukaryotic cells to prokaryotic organisms, in which the NHEJ molecular machinery is only moderately active or is entirely absent (15, 16, 23, 33) and chromosomal DNA is much more accessible (21, 34). Meanwhile, in bacterial cells, sgRNA cleavage activity can be directly related to cellular survival via the lethality that results from CRISPR/Cas9-induced DSBs with minimal perturbation from chromatin structure or DNA repair. This advantage makes it possible to prepare a large-scale, unbiased sgRNA activity dataset by designing a sgRNA library targeting every gene in the genome in bacteria, without the need to select a batch of genes with common functions, by which the bias or noise may be introduced, as has been done in mammalian cell line screening (24–29). We believe that such advantages will not only facilitate deciphering of the genome-scale sgRNA activity landscape in bacteria, but also provide more general insights into the sgRNA sequence-activity relationship based on larger and better (i.e., unbiased and with an improved signal-to-noise ratio) datasets.

In this paper, we describe a genome-wide sgRNA library consisting of roughly 70,000 members, covering both gene-coding and intergenic regions, which is comparable to ~10% of all possible sgRNA candidates (N20NGG) in the *E. coli* genome. With this library, we used a pooled screening strategy to characterize genome-wide sgRNA cleavage activity in *E. coli* by associating CRISPR/Cas9-induced DNA cleavage with cellular lethality. We observed significant sgRNA activity diversity within individual genes and across different genomic loci and thus constructed a comprehensive sgRNA activity map as a guideline for better usage of CRISPR/Cas9 genome editing tool in *E. coli*. Moreover, we found a very low correlation between our dataset and the current sgRNA activity prediction models trained from eukaryotic datasets. We therefore developed improved algorithms for sgRNA activity prediction, allowing the prediction of highly active sgRNAs in *E. coli* or, potentially, in other prokaryotic organisms. Importantly, this new model identified determinants in the sgRNA sequence for activity prediction and highlighted several key differences between wild type Cas9 and its off-target-reducing mutant.

## Results

### Design of the *E. coli* genome-wide sgRNA library and screening experiment conditions

Previously, we reported a CRISPR interference (CRISPRi) approach to perform pooled functional genomics screening in *E. coli* using a genome-wide sgRNA library consisting of 55,671 members (Wang et al, unpublished data). This CRISPRi library contains at least one sgRNA for 98.6% of 4,140 protein-coding genes and 79.8% of 178 RNA-coding genes, and most genes are targeted by 15 sgRNAs. In the current study, we repurposed this library for sgRNA cleavage activity profiling using wild-type Cas9 with nuclease activity. In addition, because the previous library includes sgRNAs targeting only the coding genome, we also designed a new sgRNA library targeting the promoter of each known operon (3,142 promoters) and ribosome-binding site (RBS) of every protein-coding gene (4,174 RBSs). These two intergenic regions are important for gene expression modulation and have attracted extensive engineering efforts (1, 35, 36). As the lengths of these intergenic regions are much shorter than the gene-coding regions, we typically designed two sgRNAs for each intergenic entry. Moreover, the sgRNAs were designed to target either of the two DNA strands in this new library, in contrast to the sgRNAs in our previous library, which bind only the nontemplate strand to maximize the CRISPRi activity. Following these guidelines, the new intergenic sgRNA library contains 10,257 members (5,559 in the promoter sub-library and 4,698 for RBSs) (Table S1 for *in silico* intergenic sgRNA library and Table S2 for promoter and RBS entries), covering 95.3% of all promoters (Figure S1) and 71.9% of all RBSs (Figure S2). Together with our previous CRISPRi library, 65,928 sgRNAs were used in this work to profile their DNA cleavage activities. This synthetic library represents a genome-wide sgRNA collection extensively covering either the gene coding or the intergenic regions of the *E. coli* genome (~10% of all sgRNAs across ~4.6 Mb *E. coli* genome assuming one sgRNA every 8 bp because of the NGG PAM requirement). Moreover, 2,000 sgRNAs without any predicted target across the *E. coli* genome were designed as an internal control in the screening experiments (Table S3). The sgRNA library was synthesized as oligonucleotides by a DNA microarray, amplified by PCR and cloned into the sgRNA expression plasmid backbone (pTargetF_lac, Wang et al, unpublished data) via Golden Gate assembly. This plasmid library was used in the following pooled screening experiments to profile sgRNA activity (Figure 1a). A log retention score (effectively the inverse of cleavage activity) for each member of the library was calculated by quantifying the representation of each sequence with and without Cas9 expression by NGS. A more negative score indicates stronger cleavage activity (Figure 1a).

**Figure 1.**
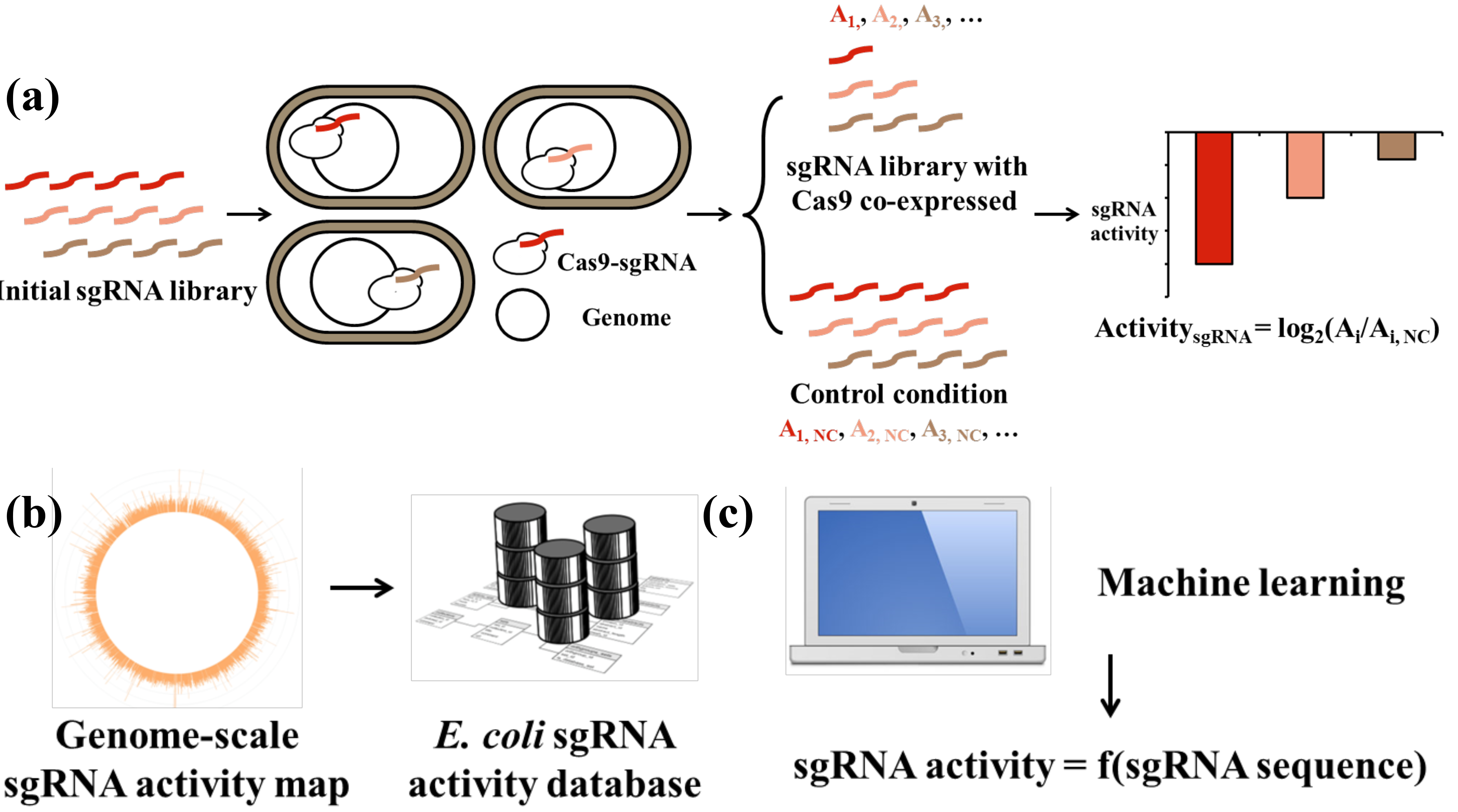
General framework combining experimental and computational approaches to depict a genome-wide sgRNA cleavage activity map in this work. (**a**) Schematic illustration of the workflow for the sgRNA activity screening experiments. The variable regions of a genome-wide sgRNA library are synthesized as oligomers on a microarray. The oligomers are subsequently amplified and cloned into an sgRNA expression vector by Golden Gate assembly. The constructed sgRNA library is transformed into *E. coli* host cells expressing Cas9 (selective condition) or dCas9 (control condition) protein. After cultivation in LB medium, the extracted sgRNA plasmids are amplified by PCR, and the abundance of each sgRNA is determined by NGS. The sgRNA activity is defined as the log_2_ change in abundance between the selective (A_i_) and control (A_i,NC_) conditions. (**b**) The obtained genome-wide sgRNA cleavage activity map can be used directly in sgRNA selection for a genome-editing project in *E. coli* (the best sgRNA for every gene, promoter and RBS encoded by *E. coli* genome). (**c**) A machine learning approach is used to shed light on the sequence-activity relationship (sgRNA activity = f (sgRNA sequence)) of sgRNAs to provide more biophysical insight into CRISPR/Cas9-based genome editing as well as to extend the sgRNA activity prediction capacity to other prokaryotic organisms.

In addition to building a comprehensive map of sgRNA cleavage activity in *E. coli* (Figure 1b), another goal of this work was to gain insight into the fundamental biophysics of the CRISPR/Cas9 system (Figure 1c). The relatively more accessible DNA substrates within bacterial cells with respect to the CRISPR/Cas9 machinery thus provide the opportunity to elucidate the inherent activity of sgRNAs based exclusively on their sequence features as well as target contexts. To this end, we characterized sgRNA cleavage activities in three different conditions (Table 1, three selective conditions). Firstly, the most widely adopted Cas9 from *Streptococcus pyogenes* was used in wild type *E. coli* strain to profile sgRNA activities. Secondly, we blocked the native DSB repair pathway of *E. coli* by deleting *recA*, which encodes a molecular sensor of DSBs and initiator of downstream homologous repair responses (18, 23). This pathway is known to play more important roles in DSB repair than NHEJ in bacteria (33). Screenings of sgRNA cleavage activities in this condition (Cas9 (Δ*recA*)) is thus expected to provide a more stable and unified baseline to dissect the underlying rules of sgRNA activities. A nuclease-dead Cas9 mutant (dCas9), which binds DNA without cleaving it, was used in wild type *E. coli* as the negative control for the abovementioned two conditions. Moreover, we also included a reengineered Cas9 derivative with improved specificity (K810A, K1003A and R1060A of Cas9) (denoted as eSpCas9) (37) in our screening experiments. For this experiment, the eSpdCas9 (K810A, K1003A and R1060A of the catalytically inactive dCas9) was used as the control condition. Table 1 summarizes the hosts and Cas9 (selective) or dCas9 (control) expression constructs for each screening experiment performed in this work and their roles in subsequent data processing to determine the sgRNA activities.

**Table 1.** Host strain and Cas9/dCas9 expression construct for each screening experiment.

| Screening experiment<sup>a</sup> | Host | Cas9/dCas9 expression vector |
| --- | --- | --- |
| Cas9 (selective) | WT<sup>b</sup> | pCas9-J23109<sup>c</sup> |
| Cas9 ($\Delta recA$) (selective) | $\Delta recA$ | pCas9-J23109 |
| dCas9 (control) | WT | pdCas9-J23109 |
| eSpCas9 (selective) | WT | peSpCas9-J23113<sup>c</sup> |
| eSpdCas9 (control) | WT | peSpdCas9-J23113 |

<sup>a</sup>Screening experiment dCas9 is the negative control for Cas9 and Cas9 ( $\Delta recA$ ); eSpdCas9 is the negative control for eSpCas9.
<sup>b</sup>WT: *E. coli* strain MCm (*E. coli* K12 MG1655 *smf::cat*, see online Methods)
<sup>c</sup>J23109 is a moderate-strength promoter, and J23113 is of weak strength.

### Quality evaluation of genome-wide sgRNA library activity profiling

To characterize the resolution of our method for differentiating among sgRNAs with diverse cleavage activities prior to screening experiments using the genome-wide library, we first applied a synthetic approach to mimic sgRNAs with a gradient of activities by introducing mismatch point mutations into the N20 region of an sgRNA targeting *yneE*. According to a previous report (11), mismatches located at the 5’ end of the protospacer region of the DNA target are better tolerated than mismatches at the 3’ end proximal to the PAM (seed region). We accordingly introduced mutations into different regions of the sgRNA to create a series of sgRNAs with diverse cleavage activities toward the same DNA substrate. In agreement with this previous knowledge, a transformation assay indeed confirmed the loss of activity as more mutations accumulated in the sgRNA N20 region that base-paired with the seed region in DNA protospacer sequence (Figure 2a). More importantly, even one mismatch mutation at the 5’ end of the sgRNA N20 region (*yneE-m1*) resulted in a 10-fold increase in survival rate with respect to wild type, which is beyond the detection limit of NGS, indicating that our method enables the discrimination of sgRNAs with only moderate activity differences.

**Figure 2.**
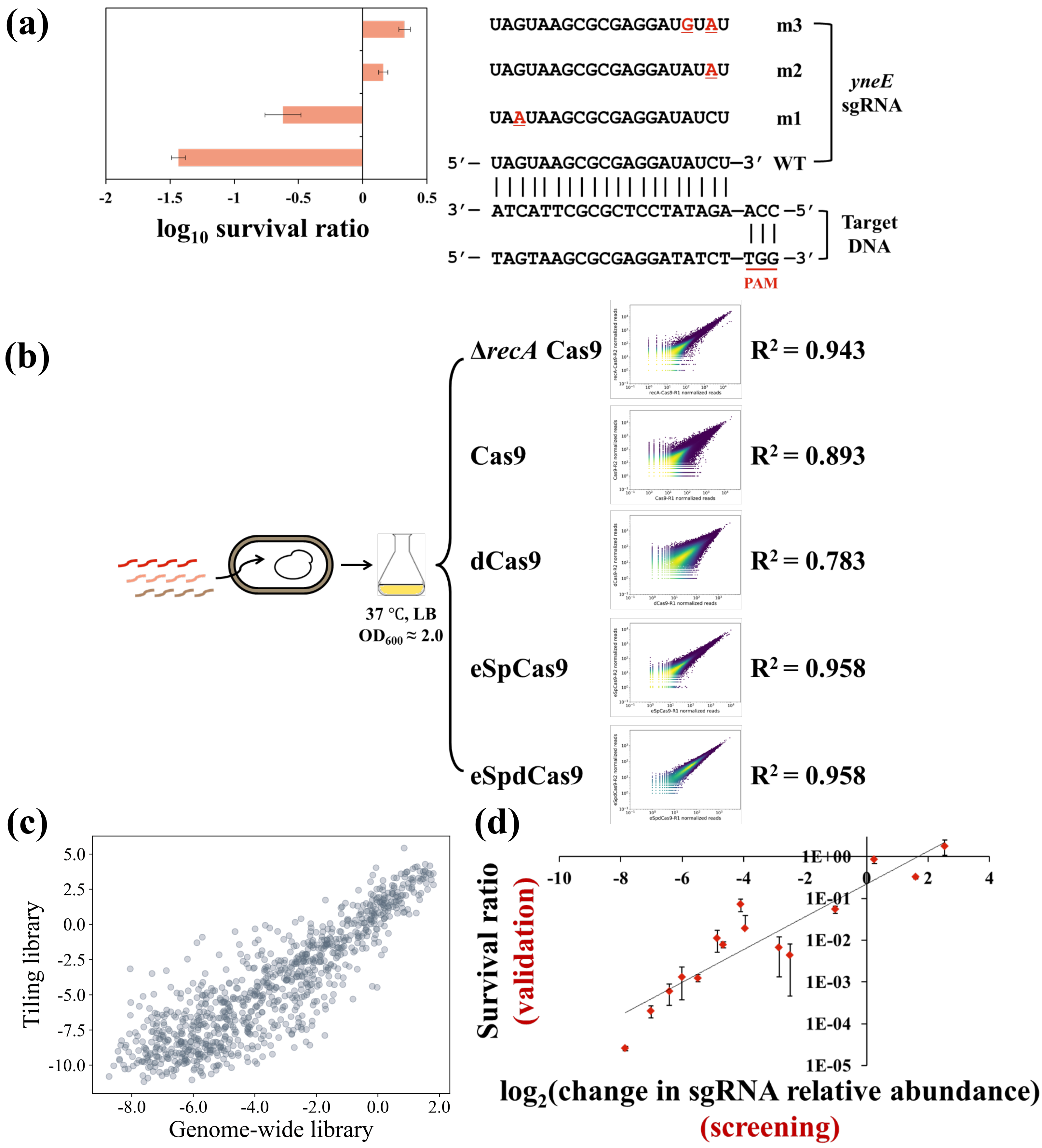
Reliability validation of screening experiments. (**a**) Different sgRNA cleavage activities are related to the cellular survival rates. Mutations were introduced at different positions in the *yneE*-WT sgRNA N20 region (*yneE*-m1, *yneE*-m2 and *yneE*-m3). These sgRNA expression plasmids were transformed into the host strains expressing Cas9 and dCas9, and the survival ratios were determined by counting the colony number after overnight cultivation on agar plates. Data represents the mean ± s.d. from n = 2 biological replicates from one experiment. (**b**) The genome-wide sgRNA cleavage activity screenings were consistent between biological replicates. Read count of each sgRNA obtained from NGS was used to compare the agreement between biological replicates (n = 2). (**c**) The results from sgRNA cleavage activity screenings were highly reproducible. One additional sgRNA library (tiling library, 3,451 members) was subjected to activity screening using the same protocol. The activity scores of 901 common members between this tiling library and the genome-wide sgRNA library obtained from relevant screening experiments are plotted against each other (R^2^ = 0.771). (**d**) The pooled sgRNA activity screening result was confirmed by cloning 15 sgRNAs individually and measuring their activities via transformation assay (colony number counting) (as in (**a**)). Data represents the mean ± s.d. of biological replicates (n = 2) from one experiment. The validation result was compared with the relative abundance changes of relevant sgRNAs obtained in high-throughput profiling (R^2^ = 0.840).

In the subsequent screening experiments, we transformed the sgRNA plasmid library into *E. coli* cells with Cas9 (selective) or dCas9 (control) expression (Table 1). The recovered culture was inoculated into fresh Luria-Bertani (LB) medium and cultivated to the exponential phase (OD_600_, ~2.0). All experiments were executed with two biological replicates. Plasmids were extracted for each culture and NGS was applied to profile each library. The consistency between replicates (Figure 2b, R^2^ > 0.78) and acceptable mapping ratio to the *in silico* library (Table S4) suggested the reliability of these experiments. To further show that the results of screenings can be reproduced in other independent experiments, we turned to a smaller tiling sgRNA library (3,451 members targeting 86 genes, Table S5). Using the same protocol as described above, we subjected this tiling library to screening for sgRNA cleavage activity with eSpCas9 in wild type *E. coli* as the selective condition. We extracted 901 sgRNAs in this tiling library, which were also included in our genome-wide sgRNA library. The comparison of activity scores for these sgRNAs between the two experiments carried out independently using different libraries suggested that our method is highly reproducible (Figure 2c, R^2^ = 0.771). To further validate results from our screenings, we selected an allelic series of sgRNAs based on their cleavage activities from the screening experiments and retested each sgRNA individually by transformation assay via colony number counting. The series consisted of 15 sgRNAs, with three sgRNAs targeting each of five genes (*ansP*-293/647/1277, *dppC*-43/637/794, *mocA*-262/294/393, *artP*-306/506/627, *araE*-595/714/1205) and with Cas9 as the selective condition in a wild-type *E. coli* host. The results showed a very good positive correlation between the screening and the validation experiments (Figure 2d, R^2^ = 0.840). Overall, these results suggested that our pooled screening method to profile sgRNA cleavage activity was very reliable and that the high-quality dataset produced accordingly could thus be used for subsequent analyses. The dataset of sgRNA cleavage activity scores obtained in this work is summarized in Tables S6 (Cas9), S7 (eSpCas9) and S8 (Cas9 (Δ*recA*)).

### The diversity of cleavage activities among sgRNAs

We first investigated the distribution of sgRNA cleavage activities obtained in the screening experiments (Figure 3a). Remarkable sgRNA activity diversity was observed for each of the three categories of conditions studied here, covering around three orders of magnitude (~10 in the log_2_ *x* axis of Figure 3a). Compared with the activity of their negative control sgRNA counterparts, the majority of sgRNAs in our library showed statistically significant cleavage activities (Figure S3, Z-score distribution). This is consistent with our previous conclusion (Wang et al, unpublished data) that ~90% of sgRNAs within the library are active for CRISPRi based gene repression. However, it should be noted that the requirements for sgRNA cleavage activity in the scenario of genome editing are much more stringent than those for CRISPRi.

**Figure 3.**
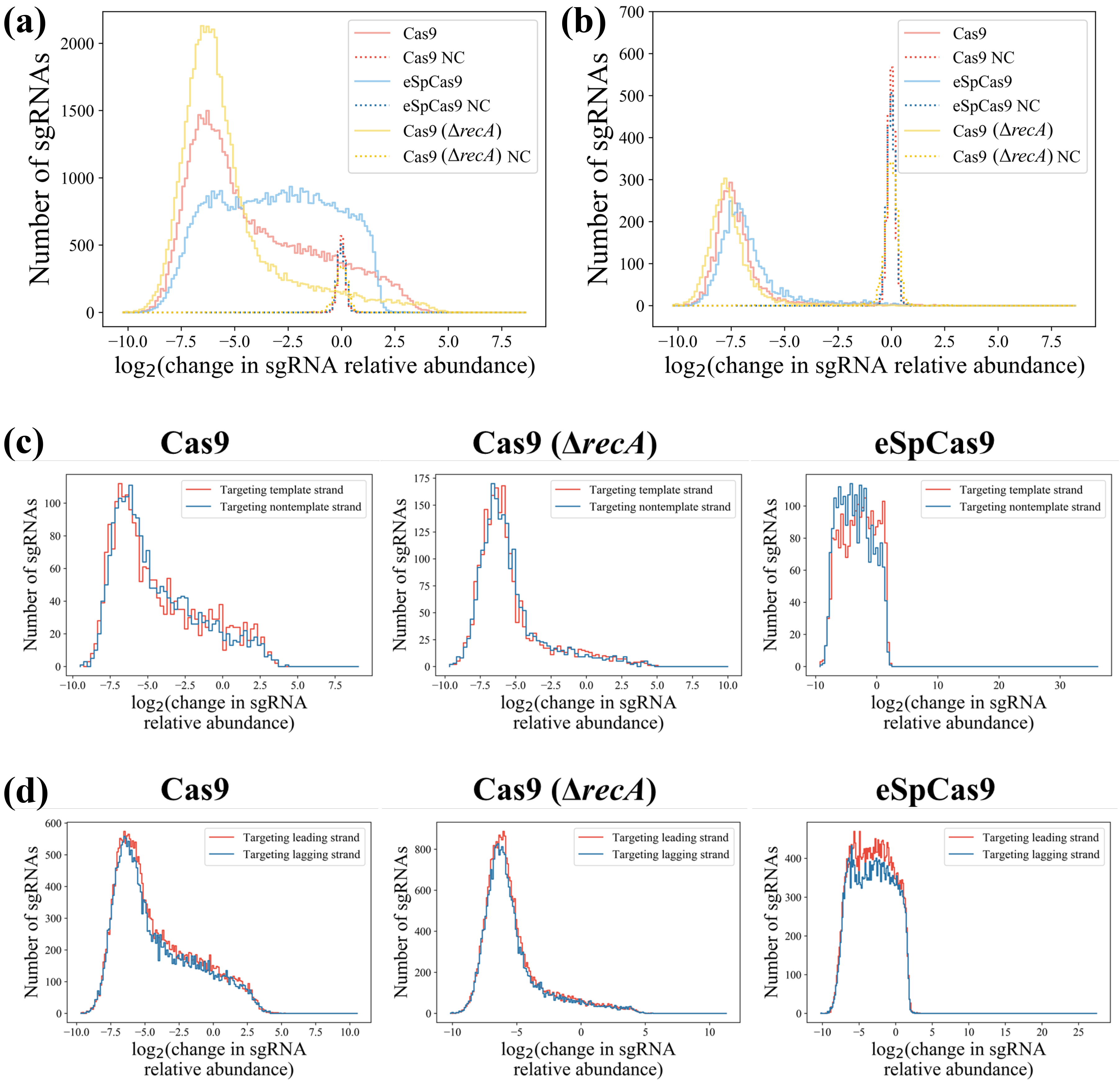
Diversity of cleavage activity among sgRNAs. (**a**) The distribution of sgRNA activity in conjunction with Cas9, eSpCas9 and Cas9 in the Δ*recA* genetic background (Cas9 (Δ*recA*)). The activity distributions of negative control sgRNAs under these three conditions are also presented as references. (**b**) Distribution of the sgRNA with the strongest cleavage activity among all sgRNAs targeting each gene (under the three conditions as in (**a**)). Only genes with at least three sgRNAs were included (4,020 genes). (**c**) Activity comparisons between sgRNAs targeting the template or nontemplate strand in the gene-coding regions. A two-tailed MW U-test was used to test for significant differences. Cas9 dataset: 2,180 vs. 2,163 (template versus nontemplate sgRNAs, respectively, here and below), *P* = 0.794; Cas9 (Δ*recA*) dataset: 2,180 vs. 2,163, *P* = 0.316; eSpCas9 dataset: 2,220 vs. 2,265, *P* = 10^−11.2^. (**d**) Activity comparisons between sgRNAs targeting the leading or lagging strand during replication across the *E. coli* chromosome. A two-tailed MW U-test was used to test for significant differences. Cas9 dataset: 27,356 vs. 25,180 (leading strand versus lagging strand sgRNAs, respectively), *P* = 0.006; Cas9 (Δ*recA*) dataset: 27,356 vs. 25,180, *P* = 0.003; eSpCas9 dataset: 29,168 vs. 26,213, *P* = 0.398.

Suppose that the upper limit of recombineering efficiency in *E. coli* is ~1% of transformants (38). In this context, among all transformants, if 1% avoid genome editing and thus survive, this results in an ~50% false positive rate (FPR). According to our comparisons between the activity scores obtained from pooled screenings and assaying colony survival after transformation for individual sgRNA (Figure 2d), we formulated a series of functions associating sgRNA cleavage activities with the FPR in genome editing based on different homologous recombination efficiencies (Figure S4). Given this relationship, it is suggested that ~40% of the sgRNAs in the Cas9 dataset (activity > −4) will lead to a FPR of 50% or higher with the optimistic assumption of 1% recombineering efficiency. Hence, we suggest that the proper selection of active sgRNAs is paramount to the success of genome editing, even in bacteria such as *E. coli* with their much more accessible chromosomal DNA than eukaryotic cells. This goal can be achieved with the help of our sgRNA activity dataset, with which we showed that at least one highly active sgRNA can be extracted for nearly all *E. coli* genes (Figure 3b). Hence, we subsequently repurposed our original sgRNA activity dataset (Table S6, Cas9) to a guideline file for genome editing to show the best sgRNA for each gene or intergenic entry (if available) encoded by the *E. coli* genome (Table S9).

We also compared the sgRNA cleavage activity profiles of the three datasets (Cas9, eSpCas9 and Cas9 (Δ*recA*)), from which the activities of the relevant nucleases can be inferred (Figure 3a). For example, the mutations in eSpCas9 are reported to increase its specificity by decreasing its stabilizing interactions with the non-target strand of the DNA substrate via eliminating the positively charged residues located within the groove between the HNH, RuvC and RNA-guided endonuclease domains (37). Our results indicated that the DNA cleavage activity of eSpCas9 is significantly weaker than that of wild-type Cas9 (Figure 3a). Thus the positively charged groove of Cas9 contributed to its endonuclease activity, although in the original report (37) the activity of eSpCas9 was not affected based on quantification of the target DNA indel mutation rate after NHEJ repair. Meanwhile, as expected, knockout of *recA* significantly increased the lethality of DSBs in the bacterial chromosome as induced by the CRISPR/Cas9 system (Figure 3a), consistent with the conclusions of previous reports(18, 23). Even though it has been suggested that genome editing can benefit from the blocking of *recA* expression (18, 39), we argue that via the rational selection of sgRNAs (Figure 3b, Table S9) the FPR of genome editing can be significantly reduced without the tradeoff of genome instability derived from blocking the inherent DNA repairing pathways in bacteria.

Most sgRNAs in our library (the CRISPRi sub-library) bind the non-template strand in the gene-coding regions, which may result in strand bias. To test whether the conclusions derived from our sgRNA library can be extended to the template strand, we used the RBS sgRNA library as a proof-of-concept because of the absence of strand bias in this library and of the location of the RBS (downstream of the transcription start site). No significant difference (Mann-Whitney (MW) U-test) was detected in the cleavage activities of sgRNAs that targeted different DNA strands for the Cas9 (*P* = 0.794) and Cas9 (Δ*recA*) (*P* = 0.316) datasets, although there was a significant difference for the eSpCas9 dataset (*P* = 10^−11.2^) (Figure 3c). One possible reason for this is that the weaker interaction of the eSpCas9-sgRNA complex with its DNA substrate (Figure 3a) may result in sensitivity to interference from the bacterial transcription machinery. Consistently, the stronger cleavage activity noted here when the nontemplate strand was targeted is consistent with a previous report that the CRISPRi system results in better gene repression when this strand is targeted (40). In spite of the strand bias noted with eSpCas9, this result indicated that there was no strand bias (transcription) in gene-coding regions for the more commonly used Cas9 in bacterial genome editing, rendering our dataset suitable for computational methods such as machine learning to predict sgRNA activities anywhere across the chromosome.

We further sought to investigate possible interactions between the CRISPR/Cas9 complex and the DNA replication machinery to identify any potential strand bias issue in this case. We divided all sgRNAs in our library into two groups based on their target DNA strand—leading strand or lagging strand during DNA replication. No significant (very weak if any) differences were found between the cleavage activities of these two groups of sgRNAs for all three datasets (Figure 3d).

### Genomic loci that are resistant to CRISPR/Cas9-induced chromosomal breaks

There are chromosomal factors such as nucleosomes that inhibit CRISPR/Cas9 genome editing in eukaryotic cells (26, 32, 41). Although it is widely accepted that bacterial DNA is much more accessible (21, 34), this issue has not yet been experimentally characterized in prokaryotic cells, in spite of the inactive genomic loci consistently observed in our datasets and the inability to modify particular genes by CRISPR/Cas9 in our experience. To this end, we used the median sgRNA cleavage activity among all sgRNAs belonging to one gene as an indicator of the average activity score of the relevant genomic region (Figure 4a). The variability in this indicator was apparent. Of particular significance is a positive skew (long right tail) in the distribution of the Cas9 dataset (Figure 4a) suggesting the existence of resistant genomic loci to CRISPR/Cas9 genome editing and thus possibly chromosomal factors that inhibit the function of the CRISPR/Cas9 complex. It is unlikely that this effect is the result of a large chromosomal deletion, because even for those genes with significant resistance to CRISPR/Cas9-induced DSBs, biologically meaningful diversity was observed for sgRNA cleavage activities within individual genes (Figure 4b).

**Figure 4.**
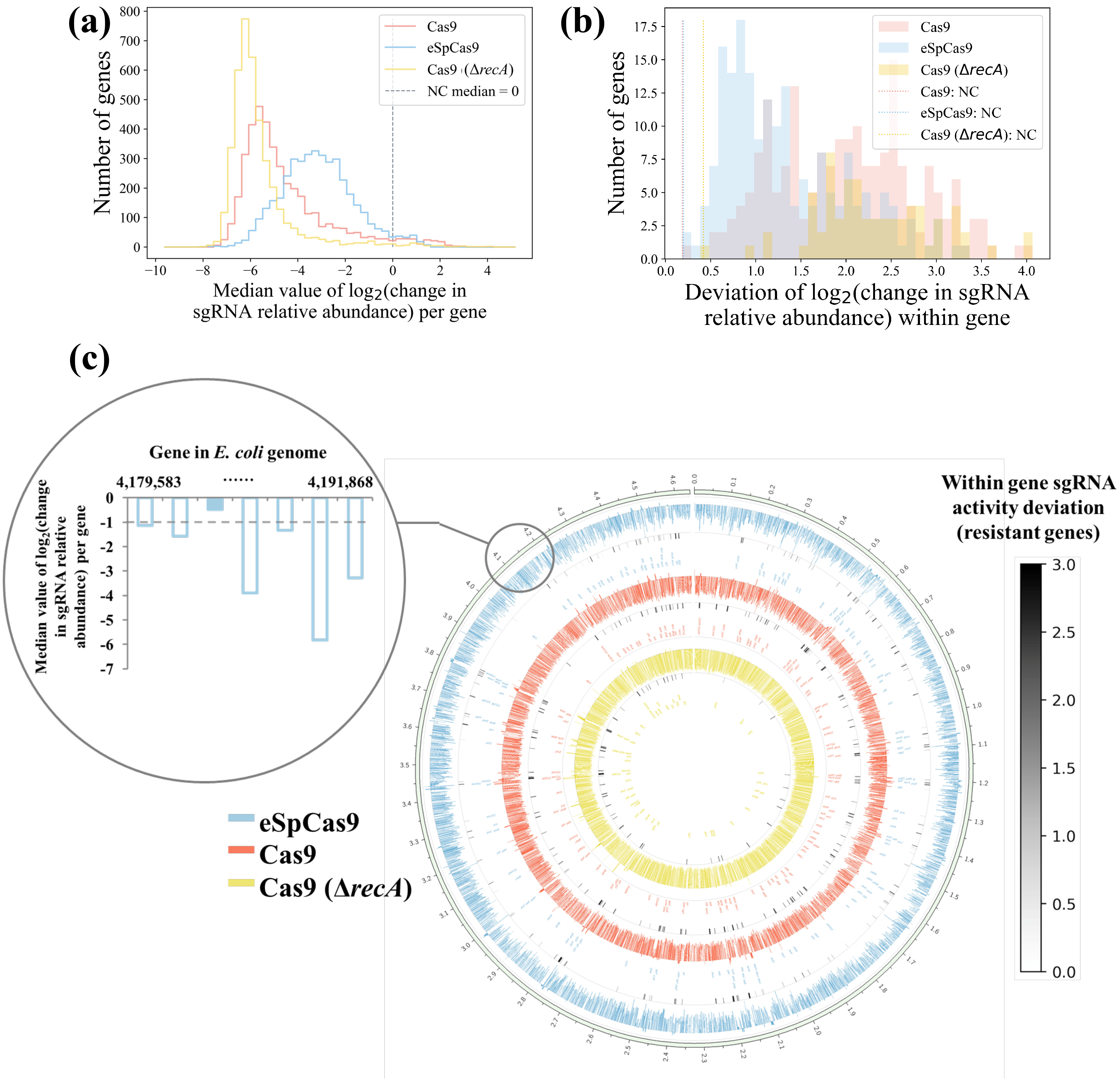
*E. coli* genome-wide landscape of resistance to CRISPR/Cas9-induced lethal DNA DSBs. (**a**) Distribution of median sgRNA activity among all sgRNAs within each gene (for each of the three conditions, Cas9, eSpCas9 and Cas9 (Δ*recA*)). The median sgRNA activity of negative control sgRNAs is zero because of the normalization step in the data processing (see Methods). (**b**) Genes with significant resistance to CRISPR/Cas9-induced DSBs were extracted ((median sgRNA activity ≥ 0 or (median sgRNA activity ≥ −1 and FPR ≥ 0. 01)) and (≥ 5 sgRNAs)), giving rise to 192, 188 and 70 genes in the Cas9, eSpCas9 and Cas9 (Δ*recA*) datasets, respectively. Shown is a distribution of deviations in the activities of sgRNAs targeting individual genes of these three sets. The experimental noise as quantified by negative control sgRNA deviations is shown as dotted lines. (**c**) Median activity among all sgRNAs belonging to each gene is plotted as a bar plot (zoom in is shown on the upper left) within each circle (Cas9, red; eSpCas9, blue; Cas9 (Δ*recA*), yellow). Genes with notable resistance to genome editing (the same threshold as in (**b**)) in each dataset (Cas9, eSpCas9 and Cas9 (Δ*recA*)) are highlighted with gene names. Essential genes in rich medium (Wang et al, unpublished data) are tagged with ‘(e)’. The heatmap (black to white) below the relevant bar plot of each circle indicates the standard deviation of the within-gene sgRNA activity for each highlighted gene. The color bar is shown on the right. A high-resolution version of this genome-wide map is accessible (https://figshare.com/s/127cecee6f9ea4e814e2) for downloading.

To illustrate the positioning of resistance to CRISPR/Cas9-induced DSBs across the chromosome, we projected the median sgRNA cleavage activity belonging to each gene encoded by the *E. coli* genome along the chromosome (Figure 4c) and highlighted those genes with poor activities. The profiles of CRISPR/Cas9-induced DSB resistance for the three datasets are consistent, especially for Cas9 and eSpCas9 (Figure 4c, three circles of white-to-black heatmaps). In contrast, some resistant genomic regions in these two datasets became vulnerable to Cas9 attack in the genetic background of Δ*recA*, suggesting that endogenous DSB repair activity sustainably mitigates the lethal effect of DSBs in a locus-dependent manner. This observation also indicates that the resistance to CRISPR/Cas9-induced DSBs of such regions in the context of Cas9 and eSpCas9 is not due to the unavailability of DNA targets via mutations such as large deletions, because all the host strains used in this work are derived from the same parental strain. This is further evidenced by the existence of within-gene sgRNA cleavage activity diversity for these resistant regions (Figure 4b and heatmap of Figure 4c). Together, our results consistently suggest that unknown chromosomal factors have an impact on the activity of CRISPR/Cas9 system, in spite of the common belief that bacterial DNA is unprotected with respect to an attack from cellular factors (21, 34). We are currently unable to associate these inactive regions with any known chromosomal factor in *E. coli*. Potential reasons for this blocking effect include DNA supercoiling state, occupation of nucleoid-associated proteins, torsional constraints of DNA, which are all suggested to impact CRISPR/Cas9 activity (28, 42) and also known to present non uniform pattern across bacterial chromosome (43–45). Given that the investigation of bacterial chromosome structure is only beginning to emerge compared with mammalian cells being extensively profiled such as in ENCODE project (46), this still needs further investigation.

### An integrated machine learning approach predicts highly active sgRNAs

As the first large-scale dataset of sgRNA cleavage activity in prokaryotes, our results make it possible to test the generalization ability of previous sgRNA activity prediction models trained by eukaryotic datasets. We adopted three widely used models (25, 28, 29) to predict the activity scores of sgRNAs in our dataset and compared those scores with our experimentally determined ones (see Methods) (Table 2, Figure S5). Two of these are machine learning models, whereas the third is based on the biophysical mechanism of CRISPR/Cas9. We found very weak correlation between our dataset and the predictions from the two machine learning models (Doench et al. and Xu et al.). In contrast, a more notable but still weakly negative correlation was observed given the predictions from the biophysical model (Farasat et al.). This result suggests that the models trained from the sgRNA activity data from mammalian cell line screenings only partially capture the patterns of sgRNA sequence-activity relationships, possibly due to the noise in the training datasets introduced via the NHEJ repair specificity (31) and the impact of dense chromatin structures (34, 41) in eukaryotic cells. Indeed, even for eukaryotic cells such as yeast, a recent study identified a different optimal window relative to the transcription start site for active sgRNA positioning in a CRISPRi system as compared with that reported for human cell lines (47). Similarly, Cui and Bikard also noted the poor prediction abilities of these models upon the cleavage activities of 13 sgRNAs in *E. coli* (23). In these lines, we sought to train the machine learning models based on our sgRNA cleavage activity dataset, aiming to extend the scope of this work to other prokaryotic organisms, as well as to elucidate the basic biophysics of the interactions between the CRISPR/Cas9 complex and DNA targets.

**Table 2.** Correlation between sgRNA activity scores from predictions based on previous mammalian-cell-line-based models and our screening experiment.

| Spearman correlation coefficient |  |  |  |
| --- | --- | --- | --- |
| Prokaryotic datasets in this work | Doench et al. (25) | Xu et al. (29) | Farasat et al. (28) |
| Cas9 dataset | 0.058 | 0.092 | -0.119 |
| eSpCas9 dataset | 0.017 | 0.052 | −0.170 |
| Cas9 $\Delta recA$ dataset | 0.012 | 0.047 | −0.068 |

We first filtered our datasets by removing sgRNAs of low quality and with multiple targets (see Methods) as well as those belonging to genes with resistance to CRISPR/Cas9 genome editing (Figure 4c). We thus established three high-quality datasets (Cas9: 44,163 sgRNAs; eSpCas9: 45,070 sgRNAs; Cas9 (Δ*recA*): 48,112 sgRNAs; summarized in Table S10) used in the subsequent work, which are the largest sgRNA on-target cleavage activity sets reported so far to the best of our knowledge (48). The absolute value of the Z-score for each sgRNA was used as the activity score. To quantitatively model sgRNA activity, we carried out a featurization process that considers the DNA target sequence of the protospacer, PAM and flanking region to convert the N20NGGN sequence into 425 binary or real number features (Figure 5a). To prevent over-fitting, we randomly separated the dataset into two subgroups with 80% of the data used as the training dataset, and the remaining 20% used to test the generalization capacity of the trained models (Figure 5a). Simple linear regression, regularized linear regression, ensemble method (gradient boosting regression tree) and an artificial neural network method (multiple layer perceptron) were used as machine learning models. We first evaluated the performance of different models using five fold cross-validation on the training set (Figure 5b). We found that the gradient boosting regression tree model was the most predictive, with Spearman correlation coefficients of 0.542, 0.682 and 0.328 for Cas9, eSpCas9 and Cas9 (Δ*recA*), respectively (Figure 5b). We reasoned that the relatively poor performance of the Cas9 (Δ*recA*) model was derived from the overwhelming selection pressure during the screening experiments (Figure 3a; see discussion in Methods), leading to the inability to discriminate between sgRNAs with strong but different activities. It is also worth noting that simple linear regression generally works quite well, with a weaker but similar correlation (Spearman correlation coefficient = 0.508, 0.624 and 0.315 for Cas9, eSpCas9 and Cas9 (Δ*recA*), respectively) with respect to that of the gradient boosting regression tree model, suggesting that the naïve linear combinations of features used here were sufficient for deciphering the sgRNA sequence-activity relationships. The trained model (fixed parameters in the final models learned from the data of training set are given in Table S11) also showed good generalization ability when predicting the unseen data in the test set, and the performance metrics were well maintained (Figure 5c). Importantly, this high predictive value was consistent across randomly selected training and test sets (data not shown). In addition, we also used the trained eSpCas9 model to carry out a prediction analysis with a dataset obtained from an independent screening experiment using the tiling sgRNA library (see Figure 2c and Methods). We applied the same filter threshold to the results of this experiment, giving rise to a high-quality set with 2,640 sgRNAs. Although 65.9% of them were not contained in the genome-wide sgRNA library, our prediction algorithm still showed good performance with a Spearman correlation coefficient value of 0.633 (Figure 5f). These results collectively confirmed that our models captured the underlying biological signals rather than fitting the data superficially. In addition, the models trained in this study (Cas9 and eSpCas9) outperformed the state-of-the-art ones in terms of Spearman correlation coefficient (25, 28, 29), possibly because of the better signal-to-noise ratio and less bias in our dataset obtained from screenings in bacteria as noted above. We propose that the algorithms reported here represent a better quantitative model about sgRNA on-target activity compared with the previous ones, at least in bacteria where it is developed.

**Figure 5.**
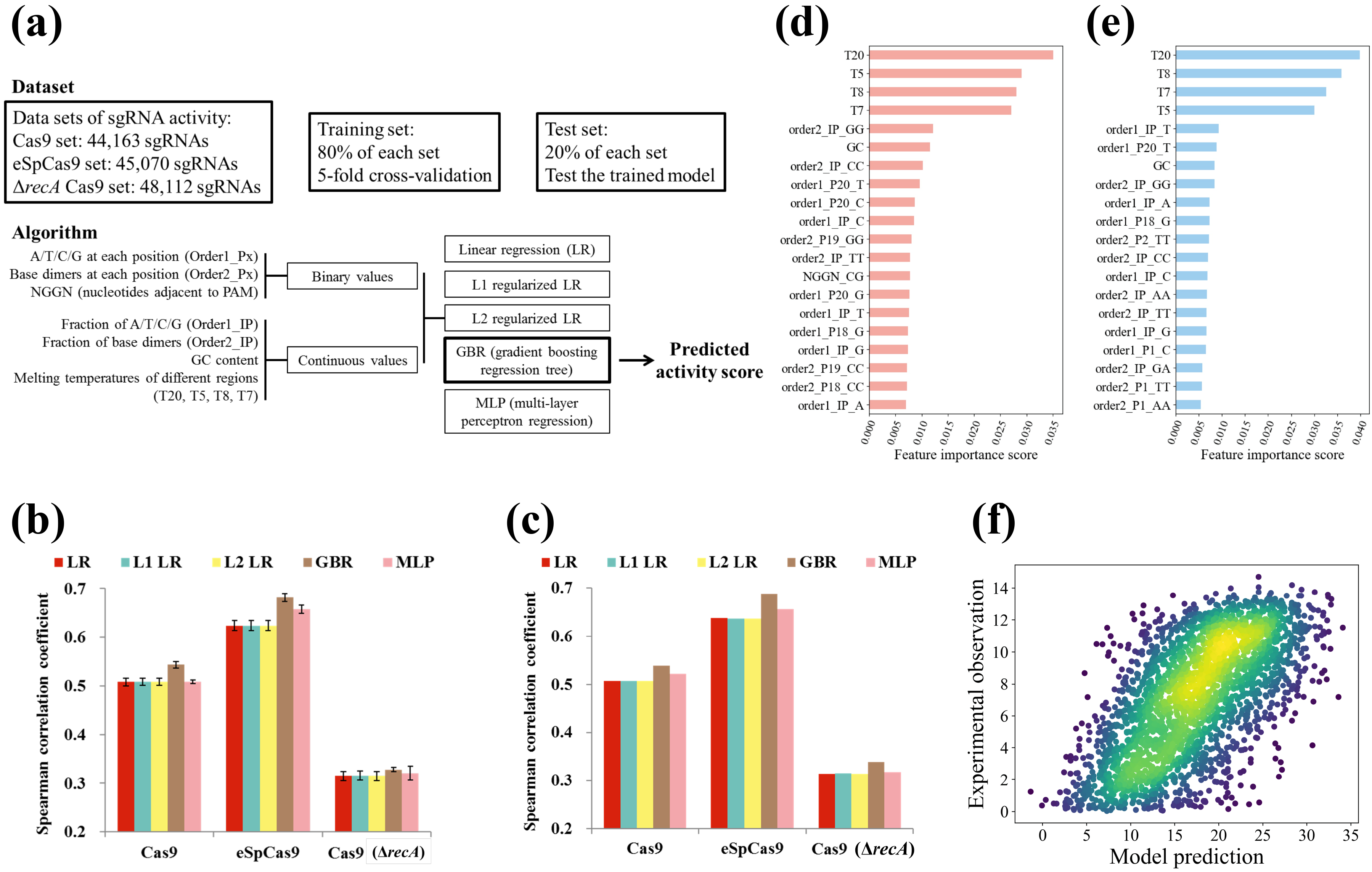
Machine learning to decipher relationships between sgRNA sequences and activities. (**a**) Schematic of machine learning dataset and algorithm. Three machine learning models (Cas9, eSpCas9 and Cas9 (Δ*recA*)) were constructed, respectively. 80% sgRNAs of the relevant dataset was used as the training set by five fold cross-validation to train the model. We reserved 20% of the sgRNAs in each set (Cas9, eSpCas9 and Cas9 (Δ*recA*)) as the test set to measure the generalization ability of each model to predict unseen data. We extracted 425 features for each sgRNA. Five varieties of machine models are trained for each dataset (Cas9, eSpCas9 and Cas9 (Δ*recA*)) and gradient boosting regression tree is found generally to perform best. (**b**) Comparison of the different models. Using fivefold cross-validation, the models were trained with the training set by 5-fold cross-validation. The bar plot shows the mean ± s.d. for the Spearman correlation coefficient between predicted and measured sgRNA activity scores (n = 5). (**c**) Comparison of the generalization ability of different varieties of models. Models were trained on the intact training set with fixed parameters optimized during cross-validation. The Spearman correlation coefficient is shown for the predicted and measured sgRNA activity scores in the test set. (**d, e**) Relative importance of features that contribute to the prediction power of the gradient boosting regression tree for the Cas9 (**d**) and eSpCas9 (**e**) model. The features of each model are sorted and the most important 20 features are shown. Results are shown from models trained on all the available data (training plus test set in (**a**)). (**f**) The generalization ability of the trained model was further validated by predicting activities from a dataset obtained from an independent sgRNA library and experiment. One additional sgRNA activity dataset (the same as that in Figure 2c) was constructed by screening the tiling library (2,640 sgRNAs passed quality control, including 901 members that were also present in the genome-wide library) using the same protocol. Predictions of sgRNA activity from this dataset based on the eSpCas9 model trained on all the available data (training plus test set in (**a**)) are plotted against experimentally obtained scores. Each point on the plots represents a unique sgRNA and color denotes the scatter density. Spearman correlation coefficient: 0.6329, *P* = 10^−294.8^.

We next analyzed which features contributed most to sgRNA activity in the gradient boosting regression tree model via Gini importance (Figure 5d for Cas9 and 5e for eSpCas9). Overall, the predicted scores were most influenced by the melting temperatures as determined by Watson-Crick base pairing. Other previously known factors that contributed to the sgRNA cleavage activity determined by our model (for both Cas9 and eSpCas9) included the inhibitory effect of extreme GC content (49) and the GG dimers (50). It is also interesting to note the differences in the profile of important features between Cas9 and eSpCas9. More well-known activity determinants of CRISPR/Cas9 activity were captured by the Cas9 model relative to the eSpCas9 model, such as the importance of a seed protospacer region proximal to the PAM site (11, 51), especially the composition of the last nucleotide (the 20^th^ nucleotide in our case (order1_P20_x)) (24); the critical role of the nucleotide immediately downstream of the NGG PAM site (NGGN_xx) (28). In the case of eSpCas9, in contrast, the seed regions were less important, and more general biophysical factors, such as the position-independent nucleotide composition (order1_IP_x or order2_IP_xx), were the predominant determinants of sgRNA on-target activity. This result suggests that the mutations may partially reprogram the recognition or subsequent interaction and cleavage function of the eSpCas9-sgRNA complex with respect to its DNA substrate, possibly because of the destabilized interaction between eSpCas9 and the non-target DNA strand resulting from the neutralization mutations to eliminate the positive charges within the inter-domain groove (37).

### Software package

To facilitate experimental biologists to use the sgRNA activity prediction models that resulted from this work, we developed an integrated Python package to convert an sgRNA sequence fasta file into activity scores. This package thus likely represents an improved alternative over existing methods optimized on datasets from mammalian cell line screenings for microbiologists and bioengineers working on bacteria. We also envision that this algorithm is useful for computational biologists to further dissect the underlying rules controlling sgRNA cleavage activities. The software can be accessed via our GitHub site (https://github.com/zhangchonglab/sgRNA-cleavage-activity-prediction.git).

## Discussion

CRISPR/Cas genome editing was elegantly demonstrated in bacteria for the first time in 2013 (10). Subsequently, a number of other groups proved the applicability of this method for a broad spectrum of prokaryotic species (12–15), including archaea (16), in which the development of tools for genetic manipulation is known to be very hard. This approach is hence regarded as a promising methodological innovation for the analysis of basic prokaryotic genetics (52) and engineering research (53, 54), such as microbial cell factory optimization or the development of a synthetic immunity arsenal to defend against pathogens. In contrast to its early optimistic expectations, the real application of CRISPR/Cas technology to microbiological or bioengineering research has lagged far behind (22, 53, 55), especially in high-throughput scenarios such as multiplex gene editing and functional genomic screening, which was previously suggested as a major advantage of CRISPR/Cas. We propose here that the sgRNA on-target activity, which has been analyzed in eukaryotic genome editing by CRISPR/Cas9 but never systematically studied in prokaryotic organisms, is a major contributor that can limit the application of CRISPR/Cas9 in bacterial gene editing. Using a comprehensive sgRNA library and pooled screening strategy, we demonstrated that sgRNA activities vary widely in *E. coli* (Figure 3a) based on sequence features (Figure 5d, e) and unknown chromosomal factors (Figure 4c), the latter of which was unexpected given the vulnerable nature of bacterial DNA. Our sgRNA activity dataset makes it possible to select optimized sgRNAs for nearly every gene and functionally important intergenic region (promoter and RBS) encoded by the *E. coli* genome (Figure 3b) and moreover to develop advanced models (Table 2 and Figure 5) to predict highly active sgRNAs not only in *E. coli* but also, potentially, in other bacteria. We believe these results should contribute to accelerating the broader and better application of promising CRISPR/Cas technology in the study of the basic biology and in the engineering of prokaryotic organisms.

In addition to these advancements in the field of bacterial genome engineering, this work also highlights the key differences in sgRNA activity profiles between Cas9 and its mutant (Figure 3a, Figure 5d, e), as well as within bacterial and mammalian hosts (Table 2). These results suggest that the underlying biophysics determining the relationship between sgRNA features and activities remains poorly understood. Notably, this work also elucidates the potential bias (hybrid of sgRNA activity and DNA repairing preference) of previous activity prediction models (25, 28, 29) trained from mammalian cell line screening data and hence raise their limitations to be applied in other CRISPR/Cas9 utilization scenario (e.g. recombination to introduce defined mutations) or to be extended to unexplored host cells, such as bacteria in this case. Hence, we suggest that more unbiased methods for high throughput sgRNA activity profiling need to be developed. Akin to the sgRNA activity datasets and the derived models reported here, such efforts should be of great value to dissect the molecular determinants of CRISPR/Cas9 genome editing activity, by which further advancing this transformative technology to realize its potential.

## Online methods

### Cell growth conditions and strain construction

In all experiments, bacteria were grown in LB medium or on LB agar plates. Cells were grown at 37 °C. Antibiotic concentrations for kanamycin and ampicillin were 50 and 100 mg/L, respectively. Molecular cloning was performed with *E. coli* DH10B as the host. *E. coli* K12 MG1655 was obtained from the ATCC (700926). The host strains used in the screening experiments were MCm and MCm Δ*recA*. MCm (Wang et al, unpublished data) was constructed by integrating a chloramphenicol expression cassette cloned from pKM154 (Addgene plasmid #13036) into the *smf* locus of wild-type *E. coli* K12 MG1655. MCm Δ*recA* was constructed by deleting the coding region of *recA* in MCm via CRISPR/Cas9 based recombineering method (17).

### Plasmid construction

The knockout of *recA* blocks DSB repair and hence boosts the lethality of the CRISPR/Cas9 system. Therefore, we chose J23113 (an Anderson promoter with weak activity) for Cas9 expression (pCas9-J23113) in host cells with the Δ*recA* genetic background (Table 1). For other cases, the medium-strength promoter J23109 was used to drive the expression of Cas9 or its derivative. To construct these plasmids, pdCas9-J23109 and pdCas9-J23113, previously described by our paper (Wang et al, unpublished data), were used as PCR templates to prepare a series of vector backbone with different promoters. The plasmid pCas (17) was used as PCR template to amplify the coding region of Cas9. These fragments were subsequently assembled via Gibson assembly to construct the intact plasmid. All sgRNA expression plasmids individually used in this work were constructed by amplifying pTargetF_lac (Wang et al, unpublished data) by PCR to alter the N20 sequence, followed by self-ligation via Gibson assembly. All the strains and plasmids used in this work are summarized in Table S12 and oligonucleotides are given in Table S13. The maps for p(d)Cas9-J23109, pCas9-J23113, peSp(d)Cas9-J23109 and representative sgRNA expression plasmids are accessible with the following hyperlinks.

https://benchling.com/s/seq-ZaTCr0hFE3U857KlIBsu

https://benchling.com/s/seq-Pk7e92yTr0X1mE9yXDeK

https://benchling.com/s/seq-23eYuaRcup6g6Ml9Dcq6

https://benchling.com/s/seq-NT6ly7Ilw3T02fpSt0iY

https://benchling.com/s/seq-sXSq0WW8RTY5lH0yRDek

https://benchling.com/s/seq-JamZWMMBAqBXhk0huc06

### Transformation assay

Cells expressing Cas9 or dCas9 were cultured overnight in LB (with kanamycin) as a seed culture followed by preparation of competent cells. Briefly, the cells were collected after growth to exponential phase (OD_600_ ≈ 0.6) by centrifugation at 8,000×g for 5 minutes at 4 °C, washed five times in ice-cold sterile water with the same condition and resuspended in 15% (v/v) glycerol (at one-sixteenth the volume of the original culture). All these operations were performed on ice. Plasmids carrying the sgRNA expression cassette (pTargetF_lac) were transformed by electroporation into the prepared competent cells expressing Cas9 or dCas9 (50 ng plasmid/100 μL competent cells). The electroporation was performed via a BTX Harvard apparatus ECM 630 High Throughput Electroporation System using an optimized parameter setting (2.1 kV, 1 kΩ, 25 μF). The transformed cells were incubated in LB medium (four times the volume of the competent cells) for 1 h at 37 °C for recovery. We streaked the resulting culture onto the LB agar plates (with kanamycin and ampicillin) automated by easySpiral Pro (Interscience). The colonies were counted after overnight cultivation. The survival ratio for each sgRNA was calculated by comparing the colony-forming units (CFU) of Cas9-expressing cells with the CFU of dCas9-expressing cells. This ratio was further normalized by determining the colony number after transformation with a negative control sgRNA plasmid to minimize the impact of differences in electroporation efficiency that were due to competent cell preparation (equation I).

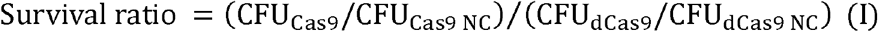

### Design and preparation of the sgRNA libraries

The contents of CRISPRi sgRNA library (Table S1) and one part of the tiling sgRNA library (see Table S5 for full list) were reported in our previous paper (Wang et al, unpublished data). We applied the same quality control threshold to design the sgRNA library targeting the promoter and RBS regions of the *E. coli* genome in this work. For the promoter sgRNA library, we downloaded the collection of *E. coli* promoters (8,594) from the RegulonDB database. Entries with overlapping regions (overlap > 1 bp) and that shared the same orientations were combined. The promoter collection was further filtered by mapping onto annotated coding genes (i.e., those that shared the same orientation and were located between 300 bp upstream of the start codon and 100 bp upstream of the stop codon), resulting in 3,146 promoters. Based on the quality control threshold described previously (Wang et al, unpublished data), every potential sgRNA (N20NGG) targeting the two strands of each promoter (40) was checked accordingly until two sgRNAs passing this threshold were extracted or the 3’ end of the promoter sequence was reached. To design the sgRNA library for RBSs throughout the *E. coli* genome, 4,140 RBS sequences for every protein-coding gene (N30 + start codon + N17, N50 in total) were extracted, and a similar procedure was applied as described above (quality control threshold and two sgRNAs per RBS). The sequences of promoter and RBS sgRNAs are summarized in Table S1, whereas library metrics and entry sequences for these two libraries are shown in Table S2. The designed sgRNAs were synthesized as oligomers on a microarray, PCR amplified and constructed as a plasmid library as described in our previous paper (Wang et al, unpublished data). We will be sharing these two sgRNA libraries through Addgene as soon as possible.

### Screening experiments

For sgRNA activity screening experiments, the single colony-derived overnight seed cultures of host strains (MCm/pCas9-J23109, MCm/peSpCas9-J23109, MCm Δ*recA*/pCas9-J23113, MCm/pdCas9-J23109 and MCm/peSpdCas9-J23109) were used to prepare competent cells as described above (transformation assay). We then mixed the library plasmids with the prepared competent cells (50 ng plasmid/100 μL competent cells) and divided the mixture into 100-μL aliquots, which were loaded into 25-well electroporation plates. The electroporation was carried out as described above using a BTX Harvard apparatus ECM 630 High Throughput Electroporation System. We typically obtained about 10^5^ colonies per well with this protocol. Two biological replicates were performed for each host strain by independent transformations. To achieve a proper coverage for the sgRNA library, we transformed 50 wells of cells for each replicate, yielding totally 10 working samples for the five host cell types two replicates each (MCm/pCas9-J23109, MCm/peSpCas9-J23109, MCm Δ*recA*/pCas9-J23113, MCm/pdCas9-J23109 and MCm/peSpdCas9-J23109). For each host, a negative control sgRNA plasmid library was also transformed using three wells of cells, which were pooled into a single independent sample, yielding five negative control libraries.

The transformed cells were incubated in LB broth (four times the volume of the competent cells) for 1 h at 37 °C for recovery. We then took a 50-μL aliquot from each culture solution, which was diluted and streaked onto LB agar plates (with kanamycin and ampicillin). After overnight incubation at 37 °C, we counted the colonies and calculated the transformation efficiency. We confirmed that each biological replicate guaranteed at least 20-fold coverage. After recovery, we inoculated the rest of each sample (replicate) into 100 mL LB broth (with kanamycin and ampicillin) in a 500-mL flask and cultivated these cells at 37 °C until an OD_600_ of ~2.0. We then took 10 mL of each resulting culture to extract plasmids using the plasmid mini kit from Omega Bio-Tek for NGS library preparation.

### NGS library preparation and sequencing

The purified plasmids were used as templates for PCR to amplify the N20 region of the genome-wide library sgRNAs (50 μL × 4 reactions per library; 50 ng template per reaction; PF/R_pTargetLacNGS_PE150 primers; KAPA HiFi HotStart polymerase (KAPA Biosystems); 95 °C 3 min, 20 cycles [98 °C, 20 s; 67.5 °C, 15 s; 72 °C, 30 s], 72 °C for 1 min). PCR conditions for tiling library is 50 μL × 4 reactions per library, 50 ng template per reaction, PF/R_pTargetLacNGS_SE50, Q5 polymerase, (NEB), 98°C 30 s, 17 cycles [98°C 10 s, 53°C 30 s, 72°C 10 s], 72°C 1 min. The sequencing library was prepared following the manufacturer’s protocol (TruSeq DNA Nano Library Prep Kit for Illumina) as described (Wang et al, unpublished data). Sequencing for the genome-wide sgRNA library was carried out using a 2 × 150 paired-end configuration and ~30 million reads were collected for each library with targeting sgRNAs and 3 million reads for negative control sgRNA libraries (Table S4). Illumina NextSeq 500 by the SE50 technique was applied for tiling sgRNA library sequencing.

### NGS data processing

Raw NGS data from each library were first combined with the relevant negative control library, resulting in 10 raw datasets (two replicates for each of the five conditions, Cas9, eSpCas9, Cas9 (Δ*recA*), dCas9 and eSpdCas9). After production of clean data by de-multiplexing and removing adaptor regions, pairs of paired-end data were merged by FLASH script (56) and those reads without detected pairs were removed. Python scripts generated in house were then used to search for the ‘GCACN20GTTT’ 28-mer in the sequencing reads (and the reverse complementary sequence), and those carrying mutations within the upstream (GCAC) or downstream (GTTT) flanking regions (4 bp each) were removed. We then mapped the extracted N20 sequences back to the *in silico* sgRNA library, via which the read count of each sgRNA in each library was determined. The mapping ratio of sequencing reads back to the *in silico* library (Table S4) for the control group was generally higher than those for selective groups, indicating the existence of selection pressure (DSB induced cell lethality) in selective groups to eliminate many sgRNAs with strong activities. For example, Cas9 (Δ*recA*) group was finally dominated by sgRNAs with synthetic errors (Table S4, ~25% mapping ratio) hence blocking CRISPR/Cas9 activity in this subpopulation. We suspected that this subpopulation is derived from the inherent error rate (~1%) in DNA oligomer synthesis, which is amplified by the selection conditions applied here. We subsequently adjusted the read counts using equation (II) (n = number of sequencing libraries) to normalize the different sequencing depths of each library. Finally, sgRNAs with <20 read counts in the plasmid library were removed to increase statistical robustness. Subsequently, the read counts for each sgRNA in the two biological replicates were averaged as the geometric mean.

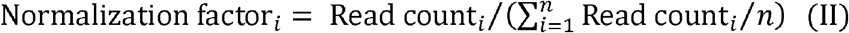

For each condition (Cas9, eSpCas9 and Cas9 (Δ*recA*)), the activity of each sgRNA was calculated via equations (III) (raw activity score) and (IV) (normalized by negative control sgRNA). Those sgRNAs with <20 reads in the control condition (dCas9 and eSpdCas9) were eliminated from the following analysis.

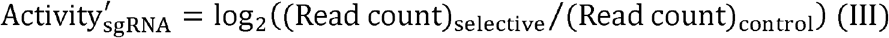

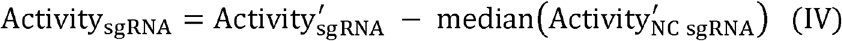

To calculate the *Z* score of each individual sgRNA, we fit the activities of all negative control sgRNAs with a normal distribution, giving rise to a value for the standard deviation (σ). The *Z* score for each sgRNA was then calculated with equation (V).

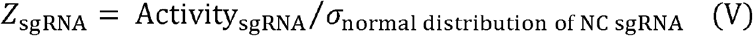

Subsequently we measured the average activity for each gene and the statistical significance in contrast to the negative control sgRNAs, to identify the genomic loci with resistance to CRISPR/Cas9-induced DSBs. Based on the activities of the sgRNAs belonging to an individual gene (including sgRNAs targeting the relevant RBS), we calculated the average based on the median of sgRNA activities and the statistical significance (FPR) via a quasi gene simulation approach (for details, see) (Wang et al, unpublished data). It should be noted that that most of the genes across the genome exhibited notable DNA cleavage activities. Hence the classical Storey-Tibshirani approach (57) for genome-wide research was not suitable here to convert the FPR value into the false discovery rate. We therefore directly used FPR values as signals to identify those resistant regions.

### Detection limit of this method

Generally, the read count for one sgRNAs was from 100 (~2^6.7^) to 1000 (~2^10^) sequencing reads (Figure 2b). This sequencing depth determined the detection limit of our method. For example, an sgRNA together with the Cas9 nuclease (selective condition) causing no doubling of the cell led to the absence (< 1 read count) of this sgRNA in the sequencing data. Hence, We reasoned that the detection limit of our method for sgRNA dropout screenings is approximately around from −7 to −10, depending on the abundance of relevant sgRNA in the initial plasmid solution for electroporation. This hypothesis was consistent with the data presented in Figure 3a (the best sgRNA gave rise to activity ~ −10). We can improve this resolution by increasing the sequencing capacity applied to each NGS library (currently 30 million reads per library). We proposed that the detection limit issue stated here was basically responsible to the poor resolution (high noise) of sgRNA activity data in Cas9 (Δ*recA*) group, because the highest selection pressure in this experiment (Table S4) resulted in the most significant sgRNA dropout (Figure 3a).

### Comparison with established models

Using the sgRNA activity datasets obtained in this work, we evaluated the performance of three previously reported activity prediction models trained based on the data from screening experiments in mammalian cell lines (Doench et al.(25); Farasat et al. (28); Xu et al. (29)). The scripts for the three sequence-activity models were downloaded (Doench et al.) or kindly provided by the relevant authors (Farasat et al. and Xu et al.), and the following commands were used to calculate an activity score for each sgRNA.

bin/SSC -l 20 -m matrix/human_mouse_CRISPR_K0_30bp.matrix -i N20NGGN7 -o output python rs2_score_calculator_v1.2.py --seq N4N20NGGN3

python Cas9_Calculator.py crRNAseq(N20) PAM(GGN) target(N20NGGN) (quickmode=False, cModelName=‘All_dataModel.mat’)

The predicted activity score for each sgRNA was compared with the experimentally determined activity value, and the Spearman correlation coefficients were calculated for each model. The high-quality sgRNA activity datasets (see below) were used here for model performance comparison rather than the full list of sgRNAs described above.

### Machine learning

#### Dataset preparation

We first carried out a filtering step to create high-quality datasets for the subsequent machine learning. We removed sgRNAs with multiple targets in the *E. coli* genome (~200 sgRNAs, for details, see Wang et al, unpublished data). Only sgRNAs targeting genes with significant cleavage activities were then kept (FPR ≤ 0.01, number of sgRNAs ≥ 5). It is noted that sgRNAs that targeted a RBS were grouped with their relevant genes, and, based on these criteria, all sgRNAs targeting promoters were removed. This filtering minimized the impact of resistant genomic loci (Figure 4c) on the quality of the dataset. The activity score for each sgRNA was calculated with equation VI, which enabled sgRNAs with better activities to have higher scores and all scores to be above or equal to zero. We thus prepared three high-quality datasets (Cas9, eSpCas9 and Cas9 (Δ*recA*); Table S10, sequence plus score) that were used in the following work.

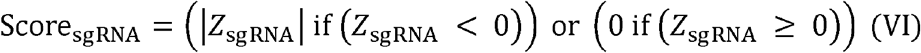

#### Featurization

We followed a featurization protocol for encoding the sgRNA sequences mainly as described by Doench et al. (25). Briefly, we used binary variables (0 or 1) to encode position-dependent base (pair) features. For example, position 1 of the 20-mer can be four different bases (A/C/T/G), each of which is encoded by four binary variables, one for each possible nucleotide. These are denoted as ‘order_1_Px’ (‘x’ denotes the position, 1–20) features, corresponding to the single base of each position. For ‘order_2_Px’ (‘x’ denotes the position, in this case 1–19) features, we looked at all adjacent dinucleotides as features, such as AA/AT/AC/AG/etc. There are 4 × 4 = 16 such base pairs, hence 16 binary variables can be used to encode one such pair at one particular position in a 20-mer. We also included position-independent features encoded by real number variables. For instance, ‘order_1_IP’ features simply mean how many A’s, etc, were in the sgRNA (20-mer), ignoring their position, as does ‘order_2_IP’. Therefore, for a 20-mer sgRNA, we obtained 80 ‘order_1_Px’, 304 (19 × 16) ‘order_2_Px’ position-specific features, 4 ‘order_1_IP’ and 16 ‘order_2_IP’ position-independent features. The two nucleotides relative to the PAM (NGGN) were also encoded, yielding 16 features, one for each NN possibility. The GC content (0–1, encoded as a real number) was computed as an additional feature. Thermodynamic features were determined via the melting temperatures of the DNA-RNA duplex using the Biopython (version 1.66) Tm_staluc function (DNA duplex version). In addition to the melting temperature of the entire 20-mer target site (‘T20’), we also included three features corresponding to the melting temperatures of three different parts of the sgRNA—the five nucleotides immediately proximal to the PAM (‘T5’), the eight nucleotides adjacent to 5’ of T5 (‘T8’), and then the seven nucleotides at the 5’ end of the 20-mer (‘T7’). We hence extracted 425 features to encode one sgRNA. These features and the sgRNA score described above were used in the subsequent machine learning.

#### Data processing for activity prediction

We first normalized the score for each sgRNA by a within-gene ranking (25) or based on the three strongest sgRNAs belonging to the gene (32). However, simple linear regression suggested there was no advantage to use these two normalized scores in contrast to the original one (Figure S6). This confirmed that our sgRNA activity screening strategy was relatively more unbiased, as compared with previous methods that associated sgRNA activities with loss-of-function phenotypes (25, 27), which makes the sgRNA activities across genes not comparable without normalization due to the differences in phenotypic effects of gene inactivation. To minimize the loss of information during data processing, we used the raw activity scores described above (equation VI) in the following work.

We used five statistical models in our experiments: (i) linear regression, (ii) L1-regularized linear regression (Lasso), (iii) L2-regularized linear regression (Ridge), (iv) gradient boosting regression tree and (v) multiple-layer perceptron. We used the scikit-learn package (0.19.0) in Python to implement each of these models. The training dataset was used for a parameter search to optimize the performance of the models by five fold cross-validation. To optimize the regularization parameter in (ii) and (iii), we searched 11 points that were evenly spaced in log space, with a minimum of 10^−5^ and a maximum of 10^5^. For the gradient boosting regression tree method, we optimized the parameters following the given order (min_samples_split, max_depth, min_samples_leaf, max_features, subsample, learning_rate and n_estimators). For multiple-layer perceptron, the regularization parameter (*alpha*) was first optimized by searching nine points that were evenly spaced in log space, with a minimum of 10^−4^ and a maximum of 10^4^. Based on the optimized regularization parameter (*alpha*), the layer topology was further optimized by searching the following combinations ([50], [100], [200], [50, 50], [100, 100], [200, 200], [50, 50, 50], [100, 100, 100], [200, 200, 200]). The optimized parameters for the five models relative to each of the three datasets (Cas9, eSpCas9 and Cas9 (Δ*recA*)) are given in Table S11.

### Statistical information, software and figure generation

Genome plots were generated using the Circos software package (58). All statistical analyses and machine learning were carried out using the SciPy (0.19.1), NumPy (1.13.1) and scikit-learn (0.19.0) Python packages. Plots were generated in Python 2.7 using the matplotlib (2.0.2) plotting libraries.

### Accession codes

Raw data of CRISPR screening for the tiling library and genome-scale library will be deposited onto the NCBI Short Read Archive as soon as possible upon publication. The software and user manual used to predict sgRNA activity can be found at https://github.com/zhangchonglab/sgRNA-cleavage-activity-prediction.git.

## Author Contributions

J.G. T.W., C.Z. and X.X. proposed the idea. T.W. designed the sgRNA library and B.L. prepared it. J.G. and T.W. performed all the experiments and carried out data processing. C.G. contributed to the data processing. J.G., T.W., C.Z. and X.X. analyzed the results. Z.X. contributed by giving critical suggestions during the project design. C.L. contributed to the technical support of machine learning. J.G. and T.W. wrote the manuscript based on discussions and contributions of all authors. C.Z. and X.X. supervised the project.

## Acknowledgements

This work was supported by the National Natural Science Foundation of China (NSFC21627812) and Tsinghua University Initiative Scientific Research Program (20161080108).

## References

1. Wang HH, Isaacs FJ, Carr P a, Sun ZZ, Xu G, Forest CR, Church GM. 2009. Programming cells by multiplex genome engineering and accelerated evolution. Nature 460:894–8.

2. Raman S, Rogers JK, Taylor ND, Church GM. 2014. Evolution-guided optimization of biosynthetic pathways. Proc Natl Acad Sci 111:17803–8.

3. Mandell DJ, Lajoie MJ, Mee MT, Takeuchi R, Kuznetsov G, Norville JE, Gregg CJ, Stoddard BL, Church GM. 2015. Biocontainment of genetically modified organisms by synthetic protein design. Nature 518:55–60.

4. Rovner AJ, Haimovich AD, Katz SR, Li Z, Grome MW, Gassaway BM, Amiram M, Patel JR, Gallagher RR, Rinehart J, Isaacs FJ. 2015. Recoded organisms engineered to depend on synthetic amino acids. Nature 518:89–93.

5. Isaacs FJ, Carr PA, Wang HH, Lajoie MJ, Sterling B, Kraal L, Tolonen AC, Gianoulis TA, Goodman DB, Reppas NB, Emig CJ, Bang D, Hwang SJ, Jewett MC, Jacobson JM, Church GM. 2011. Precise manipulation of chromosomes in vivo enables genome-wide codon replacement. Science 333:348–53.

6. Warner JR, Reeder PJ, Karimpour-Fard A, Woodruff LB a, Gill RT. 2010. Rapid profiling of a microbial genome using mixtures of barcoded oligonucleotides. Nat Biotechnol 28:856–62.

7. Pines G, Freed EF, Winkler JD, Gill RT. 2015. Bacterial Recombineering: Genome Engineering via Phage-Based Homologous Recombination. ACS Synth Biol 4:1176–1185.

8. Garst AD, Bassalo MC, Pines G, Lynch SA, Halweg-Edwards AL, Liu R, Liang L, Wang Z, Zeitoun R, Alexander WG, Gill RT. 2016. Genome-wide mapping of mutations at single-nucleotide resolution for protein, metabolic and genome engineering. Nat Biotechnol 35:48–55.

9. Cong L, Ran FA, Cox D, Lin S, Barretto R, Habib N, Hsu PD, Wu X, Jiang W, Marraffini L a, Zhang F. 2013. Multiplex genome engineering using CRISPR/Cas systems. Science 339:819–23.

10. Jiang W, Bikard D, Cox D, Zhang F, Marraffini L a. 2013. RNA-guided editing of bacterial genomes using CRISPR-Cas systems. Nat Biotechnol 31:233–9.

11. Jinek M, Chylinski K, Fonfara I, Hauer M, Doudna JA, Charpentier E. 2012. A programmable dual-RNA-guided DNA endonuclease in adaptive bacterial immunity. Science (80-) 337:816–821.

12. Oh J-H, van Pijkeren J-P. 2014. CRISPR-Cas9-assisted recombineering in Lactobacillus reuteri. Nucleic Acids Res 42:e131–e131.

13. Tong Y, Charusanti P, Zhang L, Weber T, Lee SY. 2015. CRISPR-Cas9 Based Engineering of Actinomycetal Genomes. ACS Synth Biol 4:1020–9.

14. Li H, Shen CR, Huang CH, Sung LY, Wu MY, Hu YC. 2016. CRISPR-Cas9 for the genome engineering of cyanobacteria and succinate production. Metab Eng 38:293–302.

15. Xu T, Li Y, Shi Z, Hemme CL, Li Y, Zhu Y, Van Nostrand JD, He Z, Zhou J. 2015. Efficient Genome Editing in Clostridium cellulolyticum via CRISPR-Cas9 Nickase. Appl Environ Microbiol 81:4423–31.

16. Nayak DD, Metcalf WW. 2017. Cas9-mediated genome editing in the methanogenic archaeon Methanosarcina acetivorans. Proc Natl Acad Sci U S A 114:2976–2981.

17. Jiang Y, Chen B, Duan C, Sun B, Yang J, Yang S. 2015. Multigene editing in the Escherichia coli genome using the CRISPR-Cas9 system. Appl Environ Microbiol 81:2506–14.

18. Ronda C, Pedersen LE, Sommer MOA, Nielsen AT. 2016. CRMAGE: CRISPR Optimized MAGE Recombineering. Sci Rep 6:19452.

19. Zhou Y, Zhu S, Cai C, Yuan P, Li C, Huang Y, Wei W. 2014. High-throughput screening of a CRISPR/Cas9 library for functional genomics in human cells. Nature 509:487–91.

20. Shalem O, Sanjana NE, Zhang F. 2015. High-throughput functional genomics using CRISPR-Cas9. Nat Rev Genet 16:299–311.

21. Struhl K. 1999. Fundamentally Different Logic of Gene Regulation in Eukaryotes and Prokaryotes. Cell 98:1–4.

22. Zerbini F, Zanella I, Fraccascia D, König E, Irene C, Frattini LF, Tomasi M, Fantappie L, Ganfini L, Caproni E, Parri M, Grandi A, Grandi G. 2017. Large scale validation of an efficient CRISPR/Cas-based multi gene editing protocol in Escherichia coli. Microb Cell Fact 16:68.

23. Cui L, Bikard D. 2016. Consequences of Cas9 cleavage in the chromosome of Escherichia coli. Nucleic Acids Res 44:4243–4251.

24. Doench JG, Hartenian E, Graham DB, Tothova Z, Hegde M, Smith I, Sullender M, Ebert BL, Xavier RJ, Root DE. 2014. Rational design of highly active sgRNAs for CRISPR-Cas9-mediated gene inactivation. Nat Biotechnol 32:1262–1267.

25. Doench JG, Fusi N, Sullender M, Hegde M, Vaimberg EW, Donovan KF, Smith I, Tothova Z, Wilen C, Orchard R, Virgin HW, Listgarten J, Root DE. 2016. Optimized sgRNA design to maximize activity and minimize off-target effects of CRISPR-Cas9. Nat Biotechnol 34:184–191.

26. Chari R, Mali P, Moosburner M, Church GM. 2015. Unraveling CRISPR-Cas9 genome engineering parameters via a library-on-library approach. Nat Methods 12:823–826.

27. Moreno-Mateos M a, Vejnar CE, Beaudoin J-D, Fernandez JP, Mis EK, Khokha MK, Giraldez AJ. 2015. CRISPRscan: designing highly efficient sgRNAs for CRISPR-Cas9 targeting in vivo. Nat Methods 12:982–988.

28. Farasat I, Salis HM. 2016. A Biophysical Model of CRISPR/Cas9 Activity for Rational Design of Genome Editing and Gene Regulation. PLoS Comput Biol 12:e1004724.

29. Xu H, Xiao T, Chen C-H, Li W, Meyer CA, Wu Q, Wu D, Cong L, Zhang F, Liu JS, Brown M, Liu XS. 2015. Sequence determinants of improved CRISPR sgRNA design. Genome Res 25:1147–57.

30. Lieber MR. 2008. The mechanism of human nonhomologous DNA end joining. J Biol Chem 283:1–5.

31. Chang HHY, Watanabe G, Gerodimos CA, Ochi T, Blundell TL, Jackson SP, Lieber MR. 2016. Different DNA end configurations dictate which NHEJ components are most important for joining efficiency. J Biol Chem 291:24377–24389.

32. Horlbeck MA, Gilbert LA, Villalta JE, Adamson B, Pak RA, Chen Y, Fields AP, Park CY, Corn JE, Kampmann M, Weissman JS. 2016. Compact and highly active next-generation libraries for CRISPR-mediated gene repression and activation. Elife 5:339–350.

33. Pitcher RS, Brissett NC, Doherty AJ. 2007. Nonhomologous End-Joining in Bacteria: A Microbial Perspective. Annu Rev Microbiol 61:259–282.

34. Kuzminov A. 2014. The PrecArious Prokaryotic Chromosome. J Bacteriol 196:1793–1806.

35. Bonde MT, Pedersen M, Klausen MS, Jensen SI, Wulff T, Harrison S, Nielsen AT, Herrgård MJ, Sommer MOA. 2016. Predictable tuning of protein expression in bacteria. Nat Methods 13:233–236.

36. Alper H, Fischer C, Nevoigt E, Stephanopoulos G. 2005. Tuning genetic control through promoter engineering. Proc Natl Acad Sci U S A 102:12678–83.

37. Slaymaker IM, Gao L, Zetsche B, Scott DA, Yan WX, Zhang F. 2015. Rationally engineered Cas9 nucleases with improved specificity. Science (80-) 351:84–88.

38. Costantino N, Court DL. 2003. Enhanced levels of lambda Red-mediated recombinants in mismatch repair mutants. Proc Natl Acad Sci U S A 100:15748–53.

39. Moreb EA, Hoover B, Yaseen A, Valyasevi N, Roecker Z, Menacho-Melgar R, Lynch MD. 2017. Managing the SOS Response for Enhanced CRISPR-Cas-Based Recombineering in E. coli through Transient Inhibition of Host RecA Activity. ACS Synth Biol 6:2209–2218.

40. Qi LS, Larson MH, Gilbert LA, Doudna JA, Weissman JS, Arkin AP, Lim WA. 2013. Repurposing CRISPR as an RNA-guided platform for sequence-specific control of gene expression. Cell 152:1173–83.

41. Horlbeck MA, Witkowsky LB, Guglielmi B, Replogle JM, Gilbert LA, Villalta JE, Torigoe SE, Tjian R, Weissman JS. 2016. Nucleosomes impede cas9 access to DNA in vivo and in vitro. Elife 5:e12677.

42. Räz MH, Hidaka K, Sturla SJ, Sugiyama H, Endo M. 2016. Torsional Constraints of DNA Substrates Impact Cas9 Cleavage. J Am Chem Soc 138:13842–13845.

43. Badrinarayanan A, Le TBK, Laub MT. 2015. Bacterial Chromosome Organization and Segregation. Annu Rev Cell Dev Biol 31:171–199.

44. Lal A, Dhar A, Trostel A, Kouzine F, Seshasayee AS, Adhya S. 2016. Genome scale patterns of supercoiling in a bacterial chromosome. Nat Commun 7:11055.

45. Badrinarayanan A, Reyes-Lamothe R, Uphoff S, Leake MC, Sherratt DJ, Telling A, Amit I, Lajoie BR, Sabo PJ, Dorschner MO, Sandstrom R, Bernstein B, Bender MA, Groudine M, Gnirke A, Stamatoyannopoulos J, Mirny LA, Lander ES, Dekker J. 2012. In vivo architecture and action of bacterial structural maintenance of chromosome proteins. Science 338:528–31.

46. Bernstein BE, Birney E, Dunham I, Green ED, Gunter C, Snyder M. 2012. An integrated encyclopedia of DNA elements in the human genome. Nature 489:57–74.

47. Smith JD, Suresh S, Schlecht U, Wu M, Wagih O, Peltz G, Davis RW, Steinmetz LM, Parts L, St Onge RP. 2016. Quantitative CRISPR interference screens in yeast identify chemical-genetic interactions and new rules for guide RNA design. Genome Biol 17:45.

48. Haeussler M, Schönig K, Eckert H, Eschstruth A, Mianné J, Renaud J-B, Schneider-Maunoury S, Shkumatava A, Teboul L, Kent J, Joly J-S, Concordet J-P. 2016. Evaluation of off-target and on-target scoring algorithms and integration into the guide RNA selection tool CRISPOR. Genome Biol 17:148.

49. Gilbert LA, Horlbeck MA, Adamson B, Villalta JE, Chen Y, Whitehead EH, Guimaraes C, Panning B, Ploegh HL, Bassik MC, Qi LS, Kampmann M, Weissman JS. 2014. Genome-Scale CRISPR-Mediated Control of Gene Repression and Activation. Cell 159:647–61.

50. Malina A, Cameron CJF, Robert F, Blanchette M, Dostie J, Pelletier J. 2015. PAM multiplicity marks genomic target sites as inhibitory to CRISPR-Cas9 editing. Nat Commun 6:10124.

51. Semenova E, Jore MM, Datsenko KA, Semenova A, Westra ER, Wanner B, van der Oost J, Brouns SJJ, Severinov K. 2011. Interference by clustered regularly interspaced short palindromic repeat (CRISPR) RNA is governed by a seed sequence. Proc Natl Acad Sci 108:10098–10103.

52. Peters JM, Colavin A, Shi H, Czarny TL, Larson MH, Wong S, Hawkins JS, Lu CHS, Koo B-M, Marta E, Shiver AL, Whitehead EH, Weissman JS, Brown ED, Qi LS, Huang KC, Gross CA. 2016. A Comprehensive, CRISPR-based Functional Analysis of Essential Genes in Bacteria. Cell 165:1493–506.

53. Luo ML, Leenay RT, Beisel CL. 2015. Current and future prospects for CRISPR-based tools in bacteria. Biotechnol Bioeng 113:930–943.

54. Jakočiūnas T, Jensen MK, Keasling JD. 2015. CRISPR/Cas9 advances engineering of microbial cell factories. Metab Eng 34:44–59.

55. Choi KR, Lee SY. 2016. CRISPR technologies for bacterial systems: Current achievements and future directions. Biotechnol Adv 34:1180–1209.

56. Magoč T, Salzberg SL. 2011. FLASH: Fast length adjustment of short reads to improve genome assemblies. Bioinformatics 27:2957–2963.

57. Storey JD, Tibshirani R. 2003. Statistical significance for genomewide studies. Proc Natl Acad Sci U S A 100:9440–9445.

58. Krzywinski M, Schein J, Birol I, Connors J, Gascoyne R, Horsman D, Jones SJ, Marra MA. 2009. Circos: An information aesthetic for comparative genomics. Genome Res 19:1639–1645.

